# Coral symbionts exhibit a polycistronic flavodiiron gene leading to functional proteins in photosynthesis

**DOI:** 10.1101/2021.04.03.438294

**Authors:** Ginga Shimakawa, Eiichi Shoguchi, Adrien Burlacot, Kentaro Ifuku, Yufen Che, Minoru Kumazawa, Kenya Tanaka, Shuji Nakanishi

## Abstract

Photosynthesis in cyanobacteria, green algae, and basal land plants is protected against excess reducing pressure on the photosynthetic chain by flavodiiron proteins (FLV) that dissipate photosynthetic electrons by reducing O_2_. In these organisms, the genes encoding FLV are always conserved in the form of a pair of two-type isozymes (FLVA and FLVB) that are believed to function in O_2_ photo-reduction as a heterodimer. While coral symbionts (dinoflagellates of the family Symbiodiniaceae) are the only algae to harbor FLV in photosynthetic red plastid lineage, only one gene is found in transcriptomes and its role and activity remain unknown. Here, we characterized the *FLV* genes in Symbiodiniaceae and found that its coding region is composed of tandemly repeated FLV sequences. By measuring the O_2_-dependent electron flow and P700 oxidation, we suggest that this atypical FLV is active *in vivo*. Based on the amino-acid sequence alignment and the phylogenetic analysis, we conclude that in coral symbionts, the gene pair for FLVA and FLVB have been fused to construct one coding region for a hybrid enzyme, which presumably occurred when or after both genes were inherited from basal green algae to the dinoflagellate. Immunodetection suggested the FLV polypeptide to be cleaved by a post-translational mechanism, adding it to the rare cases of polycistronic genes in eukaryotes. Our results demonstrate that FLV are active in coral symbionts with genomic arrangement that is unique to these species. The implication of these unique features on their symbiotic living environment is discussed.

## Introduction

Molecular oxygen (O_2_) is crucial for many living organisms on Earth. In particular, it is the final electron acceptor in the aerobic respiratory electron transport chain in bacteria and in the mitochondrial inner membrane of animals and plants. However, O_2_ can be converted into toxic molecules like reactive oxygen species (ROS), including the superoxide anion radical (O_2_^−^), hydrogen peroxide (H_2_O_2_), the hydroxyl radical (·OH), and singlet oxygen (^1^O_2_) (Foyer and Noctor 2020). In mitochondria, O_2_^−^ is produced in complexes I and III of the electron transport chain (Noctor et al. 2007). In plants’ thylakoid membrane of the chloroplasts and cyanobacteria, photosystem II and I (PSII and PSI) are another hot spot for generation of ROS (Krieger-Liszkay 2005; Schmitt et al. 2014; Dietz et al. 2016). During photosynthesis, O_2_^−^ is mostly generated at the PSI when the light energy is strong and excessively overcomes the demand for photosynthetic CO_2_ assimilation (Inoue et al. 1989; Kozuleva et al. 2020; Roach et al. 2015). Once generated, ROS rapidly and irreversibly cause the inactivation of PSI (Sonoike 2011). In photosynthetic organisms, the PSI reaction center chlorophyll, P700, is kept oxidized under excess light conditions by a variety of regulatory mechanisms, therefore preventing the generation of ROS in PSI (Sejima et al. 2014; Shimakawa and Miyake 2018).

In many photosynthetic organisms, flavodiiron proteins (FLV) dissipate the excess photosynthetic electrons by reducing O_2_, allowing P700 to be kept oxidized under high reductive pressure (Alboresi et al. 2019a; Shimakawa et al. 2019). Originally, photosynthetic FLV have been identified in the cyanobacterium *Synechocystis* sp. PCC 6803 (S6803) as a homologous protein to an A-type flavoprotein from anaerobic bacteria, in which the protein family is termed as FDP (Vicente et al. 2002; Helman et al. 2003). The FDP family is defined based on two functional domains: a diiron center and a flavin mononucleotide (FMN) binding motif that presumably serves the reduction of O_2_ directly to H_2_O using various electron donors (Romão et al. 2016a). In contrast to FDP in anaerobic bacteria, the FLV in oxygenic photosynthetic organisms harbors an additional unique domain that has homology with a flavin:NAD(P)H oxidoreductase (FlR) (Vicente et al. 2002). The detailed reaction mechanism of FLV is still unclear. Many mutant studies have shown that FLV are a major contributor to the electron transport from the acceptor side of PSI to O_2_ (Helman et al. 2003; Yamamoto et al. 2016; Chaux et al. 2017), which can contribute to P700 oxidation (Allahverdiyeva et al. 2013; Shimakawa et al. 2016; Helman et al. 2003). Recently, in cyanobacteria, FLV were found to reduce O_2_ by using electrons from reduced ferredoxin (or iron-sulfur clusters) *in vivo* (Sétif et al. 2020). Overall, FLV are crucial in many photosynthetic organisms to safely use O_2_ to relieve the reduction pressure on the photosynthetic electron transport chain.

Strikingly, with the exception of some species with low quality genome assemblies, FLV-containing photosynthetic organisms harbor a pair number of *FLV* genes (Allahverdiyeva et al. 2015; Alboresi et al. 2019a). In photosynthetic organisms, there are two different types of FLV isozymes: FLVA (FLV1 and FLV2) and FLVB (FLV3 and FLV4). Based on mutant studies, the presence of both FLV isozymes is necessary for the O_2_-dependent electron transport (Helman et al. 2003; Allahverdiyeva et al. 2013; Shimakawa et al. 2015; Chaux et al. 2017; Gerotto et al. 2016). Originally, bacterial FDP are known to function in the form of a head-to-tail homodimer or tetramer because the catalytic site of the diiron center is closer to that of the FMN binding site in the protein on the other side (~ 6 Å) than in that within the same monomer (~ 40 Å)(Romão et al. 2016a). In photosynthetic organisms, the loss of the FLVB isoform induces the loss of the other one (Allahverdiyeva et al. 2013; Gerotto et al. 2016; Chaux et al. 2017). Consequently, it has been proposed that FLVA and FLVB function as a heterodimer in O_2_ photoreduction (Mustila et al. 2016). However, O_2_ reduction activity has been recently reported for both FLVA and FLVB homodimers from the cyanobacterium S6803 (Brown et al. 2019) and the activity of each dimeric form *in vivo* is still unclear.

Using the FLV-specific P700 signature, the FLV-mediated electron transport has been observed in a large part of the so-called photosynthetic green plastid lineage (Ilík et al. 2017; Takagi et al. 2017). Among oxygenic photosynthetic organisms, physiological functions of FLV have been confirmed by mutant studies in cyanobacteria, green algae, liverworts, and mosses (Gerotto et al. 2016; Chaux et al. 2017; Shimakawa et al. 2017a; Alboresi et al. 2019a). The *FLV* genes are further found in ferns and gymnosperms but not in angiosperms (Alboresi et al. 2019b). On the other hand, *FLV* genes are not present in the so-called photosynthetic red plastid lineage with the notable exception of coral symbiotic dinoflagellates (family Symbiodiniaceae). Although the coral symbionts have shown large capacity of O_2_-dependent photosynthetic electron transport activity, the mechanism involved remains unknown (Roberty et al. 2014; Roberty et al. 2015).

Here, we first characterized the *FLV* gene in Symbiodiniaceae and found only one *FLV* coding sequence that is almost twice bigger than that in other photosynthetic organisms. By studying the O_2_-dependence of photosynthetic electron transport and of P700 oxidation kinetics in coral symbionts, we further show that this FLV should be involved in electron transport. Based on the amino-acid sequence alignment and phylogenetic analysis, we show that in coral symbionts the gene pair for FLVA and FLVB are fused at the transcript level although the gene products could be detected in the individual forms as well as the fused protein. The existence of a polycistronic *FLV* gene in Symbiodiniaceae suggests that both FLVA and FLVB have been required to function in protecting PSI against damage.

## Materials and Methods

### Algal and coral materials

*Cladocopium* sp. (C92, NIES-4077) was grown under a 12h/12h-light/dark cycle (23°C, fluorescent lamp, 50 μmol photons m^−2^s^−1^/21°C) in artificial sea water with an IMK medium (Nihon Pharmaceutical Co., Tokyo, Japan). Nomenclature of Symbiodiniaceae strains was defined according to (LaJeunesse et al. 2018). The coral *Acropora* sp. was obtained from Aqua Gift (Osaka, Japan) and measured within the day of purchase.

### Measurements of O_2_ and chlorophyll fluorescence

The concentration of O_2_ in the *Cladocopium* sp. culture (OD_750_ = 1.0) was monitored in an O_2_ electrode chamber (DW2/2; Hansatech Ltd, King’s Lynn, UK) simultaneously with chlorophyll fluorescence using a Dual-PAM/F (Walz, Effeltrich, Germany). Cells were illuminated with red actinic light (635 nm, 300 μmol photons m^−2^ s^−1^) at 25 °C. During the measurement, the culture was stirred with a magnetic stirrer. Pulse-modulated excitation was achieved using an LED lamp with a peak emission of 620 nm. Modulated fluorescence was measured at λ > 700 nm. The effective quantum yield of PSII, Y(II), was calculated as (F_m_′ – F′)/F_m_′ with: F_m_′, maximum fluorescence from a light-adapted cells; F’, fluorescence emission from a light-adapted cells. Short saturation flashes (10,000 μmol photons m^−2^ s^−1^, 300 ms) were applied to determine F_m_′. An anaerobic condition was prepared with glucose (5 mM), catalase (250 units mL^−1^), and glucose oxidase (5 units mL^−1^).

### Measurement of P700

The transmittance of oxidized P700 (P700^+^) was measured using a Dual-PAM/F (Walz) at room temperature (23 ± 1 °C) (Klughammer and Schreiber 1994). P700^+^kinetics during a short light (1,000 μmol photons m^−2^s^−1^) was analyzed in the coral *Acropora* sp. adapted to the darkness for 10 min. Total photo-oxidizable P700 was determined by the illumination with a short saturation flash (10,000 μmol photons m^−2^s^−1^) in the presence of 3-(3,4-dichlorophenyl)-1,1-dimethylurea (DCMU).

### Bioinformatics

Amino-acid sequence alignment of FLV isozymes was performed using Muscle (Edgar 2004) based on the coding sequences after the part of the N-terminal extension was omitted and the sequence gap was squeezed by Gap Strip/Squeeze (https://www.hiv.lanl.gov/content/sequence/GAPSTREEZE/gap.html) with 50% gap tolerance. For the alignment in Supplemental Fig. S1, the gap was not squeezed. The prediction of the motifs was performed for FLV1 and FLV3 in the cyanobacterium S6803 at GenomeNet (https://www.genome.jp) with reference to the Pfam database (Bateman et al. 2004). Protein molecular weights were predicted using tools on the ExPASy Server (https://www.expasy.org). Subcellular localization was predicted using DeepLoc (http://www.cbs.dtu.dk).

For the phylogenetic analysis, amino acid sequences of FLV isozymes were aligned by MAFFT-LINSI v7.471 (https://mafft.cbrc.jp/alignment/software)(Katoh and Standley 2013). The N terminus region corresponding to the 1-74 amino acids of FLVA in *Chlamydomonas reinhardtii* was removed in each sequence of the alignment, because the other N terminus region (75~) of *Chlamydomonas* FLVA was detected as the conserved domain NorV (accession: COG0426) by NCBI Searches for Conserved Domains (https://www.ncbi.nlm.nih.gov/Structure/cdd/wrpsb.cgi). Thereafter, the alignment was trimmed using ClipKit version 0.1 (https://github.com/JLSteenwyk/ClipKIT) with “kpic-gappy” option which keeps parsimony-informative sites and removes sites with 90 % gap (Steenwyk et al. 2020). The maximum-likelihood phylogenetic tree of FLV was inferred by IQ-TREE 2.0.7 (http://www.iqtree.org) using the best-fit model: LG+I+G4 according to BIC selected by ModelFinder stored in IQ-TREE 2.0.7 (Minh et al. 2020; Kalyaanamoorthy et al. 2017). SH-like approximate likelihood ratio test (1000 replicates), aBayes test, and ultrafast bootstrap approximation (1000 replicates) were performed (Hoang et al. 2017; Kalyaanamoorthy et al. 2017; Anisimova et al. 2011; Guindon et al. 2010). The tree was rerooted with *Escherichia coli* FDP and drawn by FigTree 1.4.4 (http://tree.bio.ed.ac.uk/software/figtree).

### Immunodetection

For the detection of FLV proteins, *Cladocopium* sp. cells were harvested from the culture (30 mL) by centrifugation, and the pellet was grounded on a mortar with liquid nitrogen. The powder was suspended in the extraction buffer containing 40 mM Tris-HCl (pH 7.4), 80 mM NaCl, 2% (w/v) sodium dodecyl sulfate (SDS), and protease inhibitor cocktail (nacalai tesque, Kyoto, Japan). After centrifugation at 13,000 × *g* for 20 min, the supernatant was incubated at 95 °C for 15 min in Laemmli SDS sample buffer and then centrifuged again. Protein concentration was determined using a Qubit fluorometer (Thermo Fisher Scientific, Waltham, MA, USA). A portion of the resulting supernatant (40 μg protein) was analyzed by SDS-polyacrylamide gel electrophoresis. After electrophoresis, the proteins were electrotransferred onto a polyvinylidene fluoride membrane and detected by an antibody raised against *Chlamydomonas* FLVA recombinant protein (Chaux et al. 2017).

## Results

### Gene structure of FLV in coral symbionts

Whereas in most of cyanobacteria, green algae, and basal land plants, FLVA and FLVB isozymes are located at different places in the genome, some cyanobacteria species such as and FLVB respectively) as neighbors in the genome (Fig. 1). Some other cyanobacteria species such as S6803 have an additional pair of FLVA and FLVB, i.e., FLV2 and FLV4, resulting in the *FLV4-2* operon (Fig. 1), which is expressed in response to CO_2_ limitation (Shimakawa et al. 2015; Santana-Sanchez et al. 2019). A CO_2_ limitation-inducible protein (ColA), a unique protein to some cyanobacteria, is also included in the *FLV4-2* operon except for *Synechococcus* sp. PCC 7002 (S7002) having ColA alone (Shimakawa et al. 2017b; Bersanini et al. 2017). We searched for analogous genes to cyanobacterial FLV in the transcriptomic data of three strains *Synechococcus elongatus* of Symbiodiniaceae, *Symbiodinium tridacnidorum*, *Cladocopium* sp. (C92), and *Durusdinium trenchii* (Shoguchi et al. 2018; Shoguchi et al. 2020). In all the three Symbiodiniaceae strains, only one *FLV* homolog could be found (Fig. 1). In order to further analyze these atypical *FLV* genes, we looked for classical FLV protein motifs in the predicted Symbiodiniaceae FLV proteins. Two independent FlR-like domains were determined one by one respectively in upstream and downstream regions in the coding sequences, which suggested that these large *FLV* genes are composed of double coding regions of common FLV tandemly repeated. We conclude that FLV in Symbiodiniaceae are probably combined in a polycistronic loci, a rare feature in eukaryotes.

**Fig. 1.**
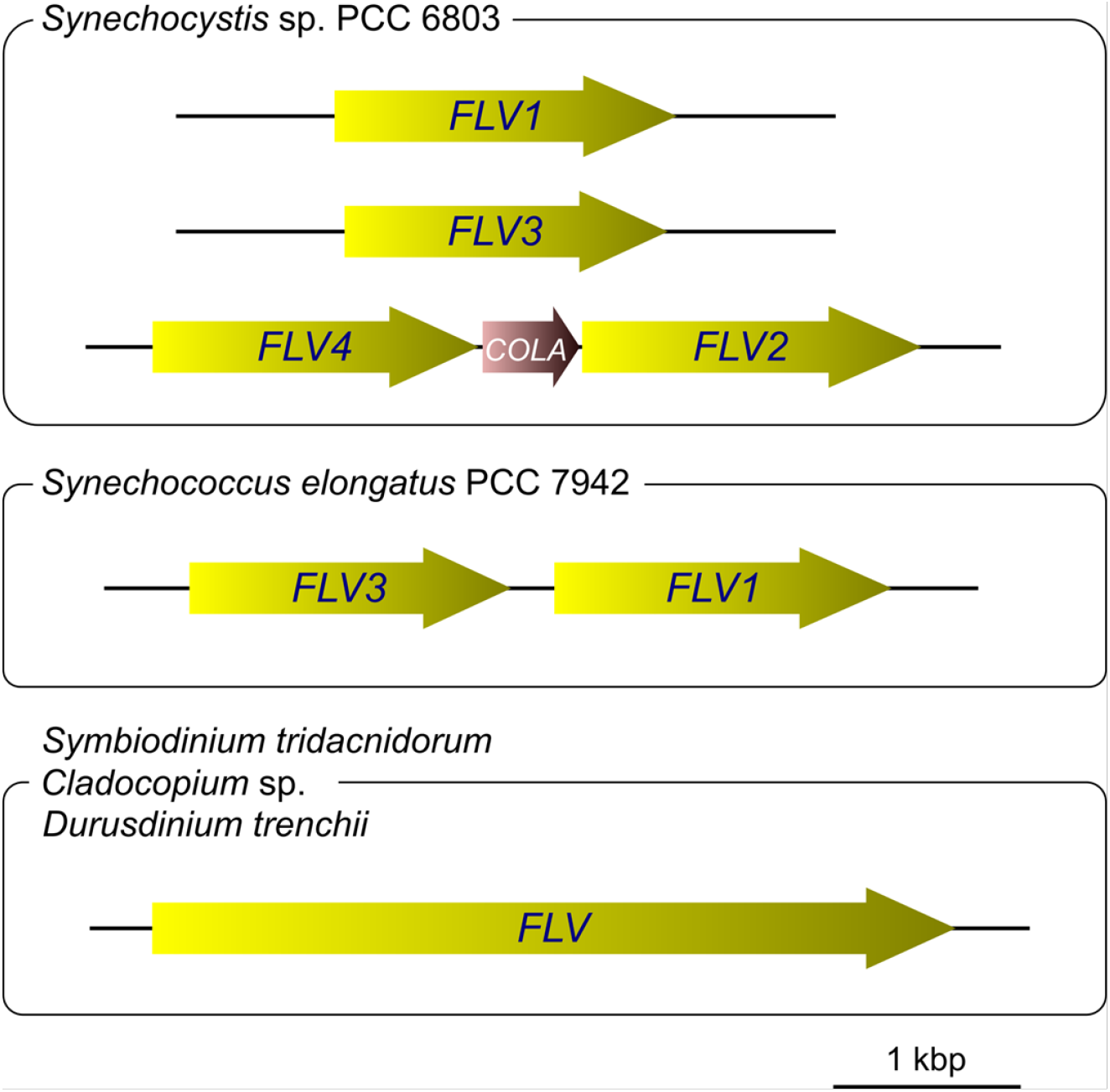
Organization of genes encoding flavodiiron proteins (*FLV*) in cyanobacteria *Synechocystis* sp. PCC 6803 and *Synechococcus elongatus* PCC 7942, and the coral symbiont *Cladocopium* sp. *COLA*, a CO_2_ limitation-inducible protein conserved in limited cyanobacteria species.

### O_2_-dependent electron transport in the coral symbiont

To assess the potential functioning of FLV in Symbiodiniaceae, we measured net photosynthetic O_2_ evolution rate simultaneously with the effective quantum yield of PSII, Y(II), during the induction phase of photosynthesis in dark-adapted cells of the Symbiodiniaceae *Cladocopium* sp. Y(II) is proportional to the activity of PSII, and therefore reflects electron flow towards CO_2_, O_2_ or any other acceptor (Shimakawa et al. 2017a; Genty et al. 1989). However, contrarily to other photosynthetic electron flux a photosynthetic electron flow towards O_2_ should not contribute to net O_2_ evolution. Thus, any discrepancy between Y(II) and net O_2_ evolution kinetics arises from an electron transport to O_2_. In *Cladocopium* sp., whereas the O_2_ evolution rate gradually increased and took approximately 3 min to reach the maximum value, Y(II) was almost at plateau already 15 s after starting illumination (Fig. 2). Furthermore, the large part of the extra Y(II) during photosynthesis induction was suppressed in the absence of O_2_ (Fig. 2). From these results we conclude that *Cladocopium* sp. possesses a strong capacity of O_2_- dependent electron transport, in agreement with previous reports (Roberty et al. 2014; Roberty et al. 2015).

**Fig. 2.**
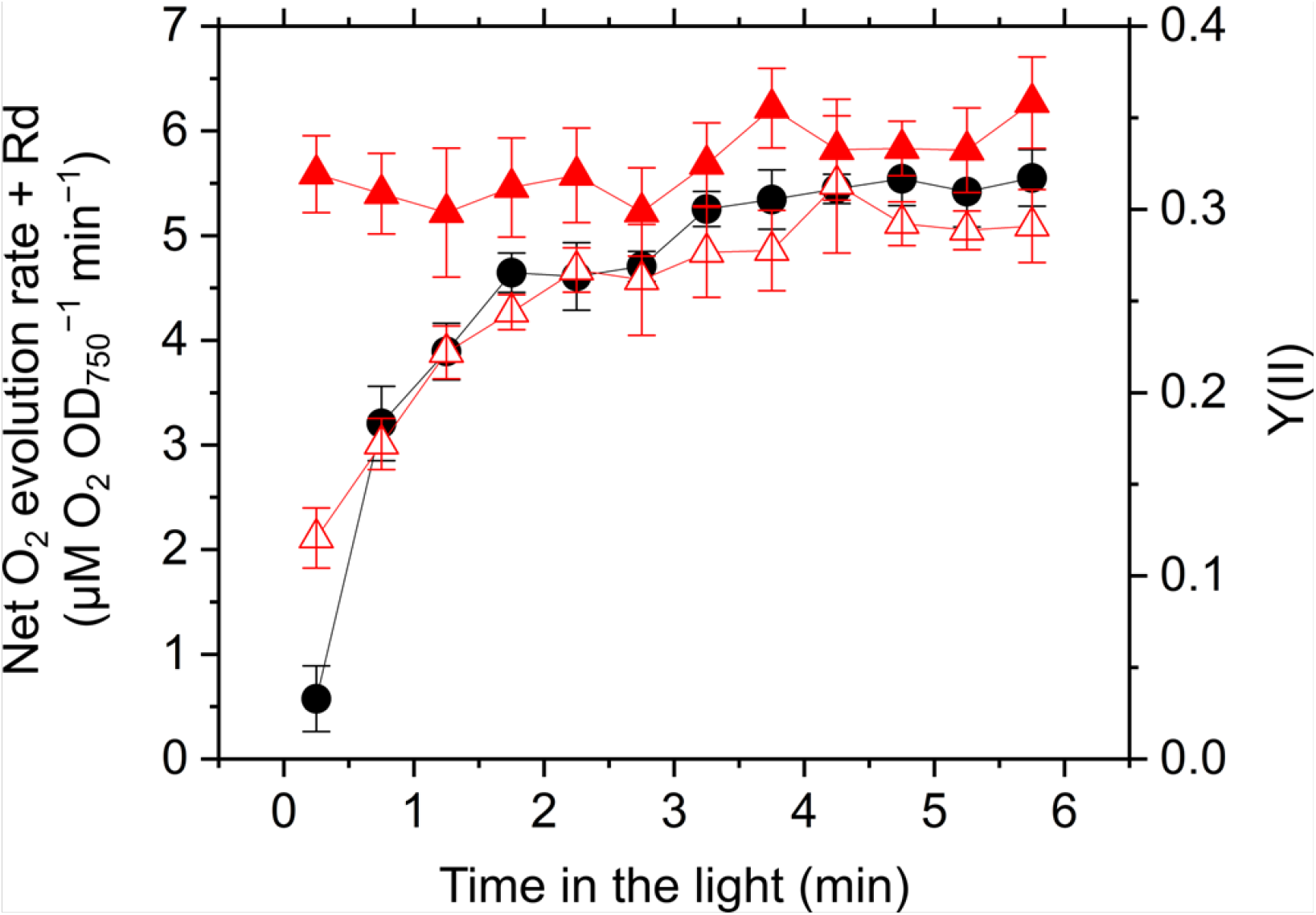
Time courses of photosynthetic O_2_ evolution rate (black circles) and effective quantum yield of PSII, Y(II) (red triangles), in the induction phase of photosynthesis in the coral symbiont *Cladocopium* sp. Actinic light (300 μmol photons m^−2^ s^−1^) was turned on at time zero. Data are represented as the mean ± SD of three independent measurements (biological replicates). Y(II) were measured in the presence (closed triangles) and absence of O_2_ (open triangles).

### P700 oxidation in the coral symbiont

To evaluate whether the observed O_2_-dependent electron transport could be ascribed to FLV activity, we evaluated the P700^+^ kinetics during the illumination with a short saturation light in the coral *Acropora* sp., because a rapid induction of P700 oxidation is characteristic of a FLV-dependent electron sink at the acceptor side of PSI (Ilík et al. 2017; Takagi et al. 2017). In the coral *Acropora* sp., a large part of P700 was kept oxidized during illumination in air (Fig. 3), with a kinetic typical to FLV activity. In addition, *Acropora* sp. could not keep P700 oxidized when aerated with Argon (Ar) to eliminate O_2_ (Fig. 3), similar to cyanobacteria, green algae, and basal land plants (Shimakawa et al. 2019; Takagi et al. 2017). The rapid induction of P700 oxidation was restored where Ar gas was replaced with ambient air (Fig. 3). We conclude from these experiments that FLV most probably mediate O_2_ photoreduction in the coral symbiont.

**Fig. 3.**
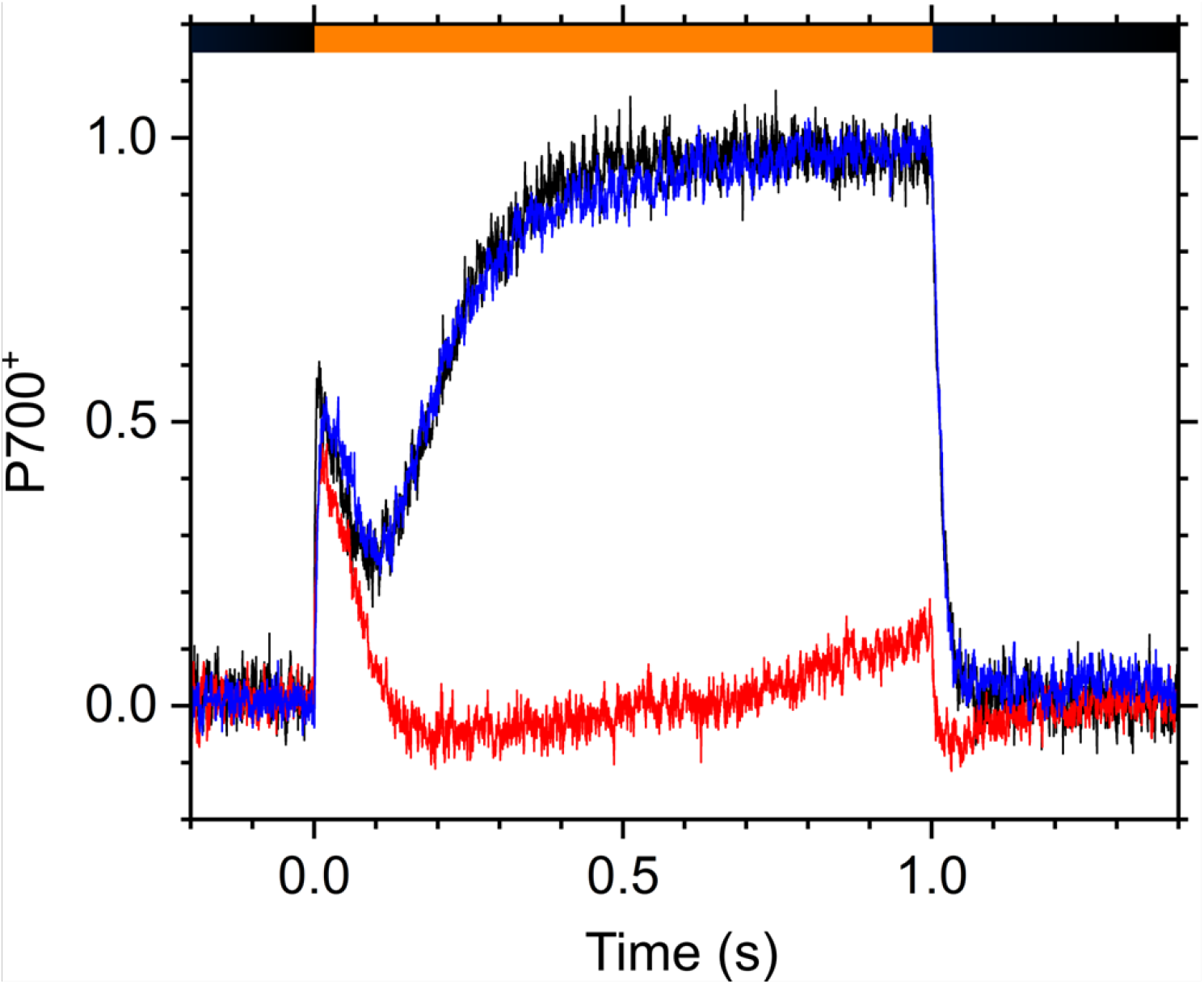
Kinetics of oxidized P700 (P700^+^) in the illumination with a short-pulse light (1,000 μmol photons m^−2^s^−1^, 1 s, orange bar) in the coral *Acropora* sp. First, experiments were performed under ambient air (black). Thereafter the gas phase was aerated with Ar (red) and then with ambient air again (blue). Relative P700^+^amount is normalized by the maximum oxidation level of P700 in the presence of 3-(3,4-dichlorophenyl)-1,1-dimethylurea (i.e., total oxidizable P700) as 1.0. Representative traces of the measurements for three different parts of the coral are shown.

### Primary structure of FLV in Symbiodiniaceae

We separated the coding regions into two parts “up” and “down (dn)” to compare the amino acid sequences with those of FLV in three cyanobacteria species S6803, S7942, and S7002, the green algae *Ostreococcus tauri*, *Micromonas* sp. RCC299, *Chlamydomonas reinhardtii*, and *Coccomyxa subellipsoidea*, the liverwort *Marchantia polymorpha*, the moss *Physcomitrella patens*, and the fern *Selaginella moellendorffii*. Whereas the amino acid sequences of three motifs specific to FLV (i.e., the diiron centre, FMN binding, and FlR-like) were similar among all these sequences, one insertion was found specific to the FLVA isozymes in *Chlamydomonas*, *Coccomyxa*, *Marchantia*, *Physcomitrella*, and *Selaginella* (green shading in Supplemental Fig. S1), and the other insertion was only specific to the upstream regions of FLV in Symbiodiniaceae (yellow shading in Supplemental Fig. S1). All the ligands binding to iron ions, commonly observed in FLVB but not in FLVA (Romão et al. 2016b), are conserved in the downstream regions but not in the upstream ones of Symbiodiniaceae FLV (Fig. 4). It has been found that FLV1 and FLV3 interact with ferredoxin in S6803 (Hanke et al. 2011), and these FLV utilize reduced ferredoxin (or iron-sulfur clusters) as the electron donor *in vivo* in S6803 (Sétif et al. 2020). Here, we found that in the motif on the electron donor side of the protein, i.e., FlR-like domain (Romão et al. 2016a), several basic amino acid residues, arginine (Arg, R) and lysine (Lys, K), are conserved in most organisms including Symbiodiniaceae (Fig. 4). This supports the Symbiodiniaceae FLV interaction with the acidic protein ferredoxin (Hase et al. 2006) and probable capacity to use ferredoxin as the electron donor, like cyanobacterial one (Sétif et al. 2020). From these data, we conclude that the *FLV* gene in Symbiodiniaceae is composed of an atypical gene fusion of eukaryotic FLVA and FLVB.

**Fig. 4.**
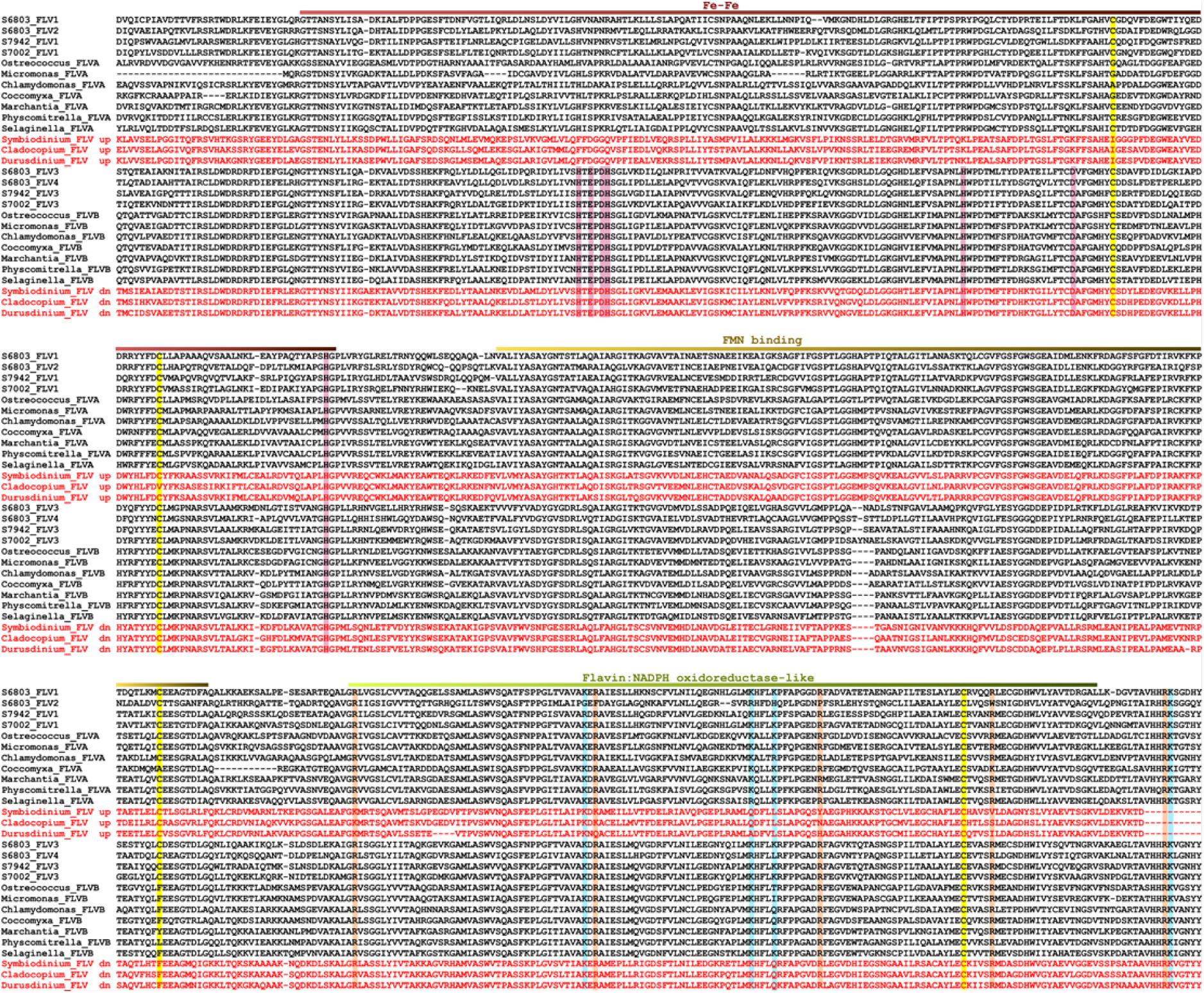
Amino acid sequence alignment of flavodiiron proteins (FLV) in cyanobacteria *Synechocystis* sp. PCC 6803 (S6803), *Synechococcus elongatus* PCC 7942 (S7942), and *Synechococcus* sp. PCC 7002 (S7002), green algae *Ostreococcus tauri*, *Micromonas* sp. RCC299, *Chlamydomonas reinhardtii*, and *Coccomyxa subellipsoidea*, the liverwort *Marchantia polymorpha*, the moss *Physcomitrella patens*, the fern *Selaginella moellendorffii*, and dinoflagellates Symbiodiniaceae (coral symbionts) clade A3 (*Symbiodinium tridacnidorum*), C (*Cladocopium* sp.) and D (*Durusdinium trenchii*). The sequences in Symbiodiniaceae were analyzed in the separated forms to upstream (up) and downstream (dn) regions. Sequence gaps in the alignment were stripped (see “Materials and Methods”). Brown, yellow, and green bars indicate diiron centre (Fe-Fe), flavin mononucleotide (FMN) binding domain, and flavin:NAD(P)H oxidoreductase-like motif, respectively. Purple shadings show ligands binding iron ions. Orange and blue shadings indicate the conserved arginine and lysine residues in the flavin:NAD(P)H oxidoreductase-like motif. Cysteine residues are indicated by yellow shadings.

### Evolutionary relationship of Symbiodiniaceae FLV with those in the other photosynthetic organisms

To get a glimpse into the evolutionary history of this atypical FLV fusion protein, we performed phylogenetic analysis using amino acid sequences from common FLV of cyanobacteria, green algae, and basal land plants (Fig. 5). We note that FLV in photosynthetic organisms can be categorized separately from FDP in other bacteria because of the additional FlR domain (Romão et al. 2016a). Therefore, we used the FDP of *Escherichia coli* as an outgroup. In the phylogenetic tree, two independent groups showing FLVA and FLVB were clearly determined (Fig. 5), which is in agreement with the previously reported phylogeny of FLV in photosynthetic green plastid lineage (Alboresi et al. 2019b). Interestingly, the two separated parts of Symbiodiniaceae FLV were obviously categorized into FLVA and FLVB respectively (Fig. 5), confirming that the large *FLV* genes in the coral symbionts tandemly encode two types of FLV isozymes. Note here that from the phylogenetic tree in this study, the upstream FLVA part of Symbiodiniaceae FLV was mutated larger than the FLVB part when they were inherited to the dinoflagellate (Fig. 5).

**Fig. 5.**
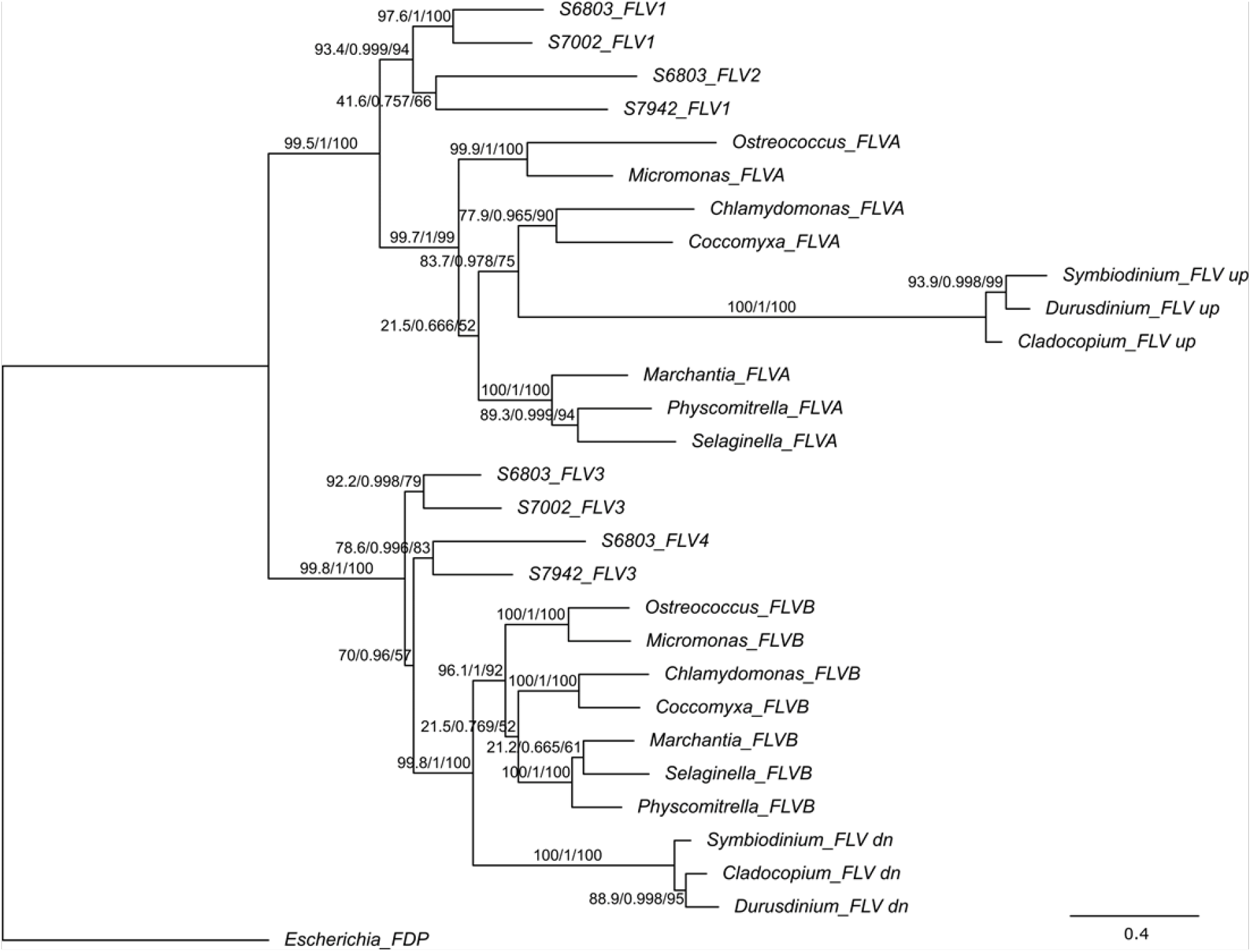
Maximum likelihood phylogenetic tree of the evolutionary relationship between flavodiiron proteins (FLV or FDP) in cyanobacteria *Synechocystis* sp. PCC 6803 (S6803), *Synechococcus elongatus* PCC 7942 (S7942), and *Synechococcus* sp. PCC 7002 (S7002), green algae *Ostreococcus tauri*, *Micromonas* sp. RCC299, *Chlamydomonas reinhardtii*, and *Coccomyxa subellipsoidea*, the liverwort *Marchantia polymorpha*, the moss *Physcomitrella patens*, the fern *Selaginella moellendorffii*, and dinoflagellates Symbiodiniaceae (coral symbionts) clade A3 (*Symbiodinium tridacnidorum*), C (*Cladocopium* sp.) and D (*Durusdinium trenchii*). The sequences in Symbiodiniaceae were analyzed in the separated forms to upstream (up) and downstream (dn) regions. The tree was rerooted with FDP of *Escherichia coli*. Branch lengths correspond to the evolutionary distances. Values of SH-aLRT support (%), aBayes support, and ultrafast bootstrap support (%) are indicated at nodes.

### Expression of FLV in the coral symbiont

To assess the protein form of FLV functioning in the coral symbionts, we detected the FLV protein in the crude soluble extract from *Cladocopium* sp. using a FLVA antibody. We detected a protein band around 85 kDa (Fig. 6), which is almost equal to the predicted size of the upstream part of the FLV corresponding to FLVA. Another band around 75 kDa was seen that matched the downstream part of the FLV corresponding to FLVB. Although weaker, we could detect a band around 150 kDa, likely corresponding to the predicted size of whole hybrid FLV protein. We conclude from this experiment that Symbiodiniaceae FLV is present in both cleaved and uncleaved forms in the cells. These results further suggested that the hybrid FLV protein is cleaved by a post-translational system and that Symbiodiniaceae *FLV* is a polycistronic gene, a rare feature in eukaryotes.

**Fig. 6.**
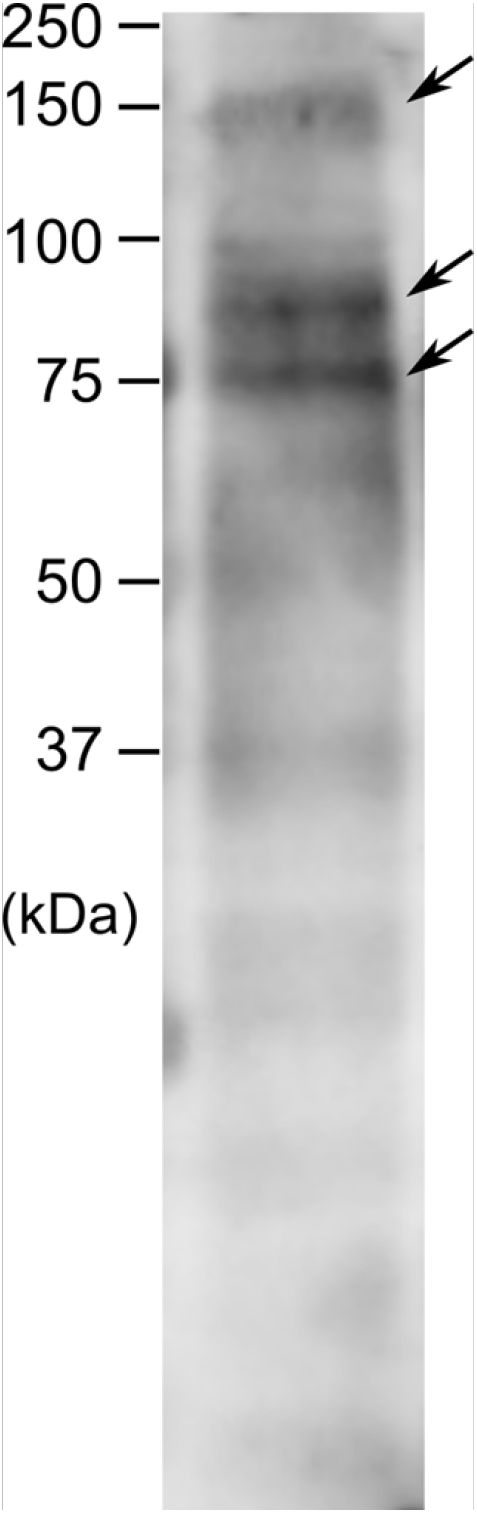
Detection of flavodiiron proteins (FLV) in a soluble crude extract of *Cladocopium* sp. Proteins (40 μg) were loaded and analyzed by SDS-PAGE, and thereafter immunoblotted with the antibody raised against FLVA. We assumed that FLV proteins were detected in the fusion (upper band) and cleaved forms (the upstream FLVA, middle band; the downstream FLVB, lower band).

## Discussion

While the flavodiiron proteins encoding genes were previously found in the coral symbionts Symbiodiniaceae genomes, it has remained unknown whether FLV were active in Symbiodiniaceae and what gene structure they exhibited. In this work, we have shown that the *FLV* gene sequences of Symbiodiniaceae are atypically composed of two classical *FLV* sequences tandemly repeated (Fig. 1). The upstream and downstream regions show strong similarity to the two photosynthetic FLV groups i.e., FLVA and FLVB (Fig. 4). From the evolutionary relationship among FLVA and FLVB isozymes in photosynthetic organisms (Fig. 5), it is highly likely that the different types of *FLVA* and *FLVB* genes have been fused in Symbiodiniaceae to result in the hybrid *FLV* gene after the primary *FLV* gene duplication common to all photosynthetic organisms. We have further shown that the FLV protein was probably cleaved into both FLVA and FLVB after translation, making it one of the rare examples of polycistronic eukaryotic gene. We have shown that the Symbiodiniaceae *Cladocopium* sp. has a strong capacity for O_2_ photoreduction comparable to CO_2_ assimilation (Roberty et al. 2014) during the first minutes of onset of light, which is similar to the FLV activity in cyanobacteria and green algae (Helman et al. 2003; Burlacot et al. 2018; Chaux et al. 2017; Allahverdiyeva et al. 2013). This important O_2_-photoreduction could be attributed to various mechanisms among such as photorespiration, FLV-mediated O_2_ photoreduction, the Mehler reaction, plastid terminal oxidase activity, and mitochondrial respiration (Curien et al. 2016). Although the lack of available genetic engineering techniques in *Cladocopium* sp. limits a dissection of the contribution of these mechanisms, the O_2_-dependent electron transport shown in this work is typical for the FLV-mediated O_2_ photoreduction measured in cyanobacteria, green algae, and basal land plants (Helman et al. 2003; Burlacot et al. 2018; Chaux et al. 2017; Santana-Sanchez et al. 2019; Shimakawa et al. 2017a). Further, *Cladocopium* sp. rapidly induced P700 oxidation in response to the illumination with actinic light, in an O_2_-dependent manner (Fig. 3), which is a typical physiological response observed in photosynthetic organisms harboring FLV (Shimakawa et al. 2019; Takagi et al. 2017; Ilík et al. 2017). We therefore propose that the polycistronic-expressed FLV is responsible for the strong O_2_-dependent electron transport during induction of photosynthesis in coral symbionts.

In cyanobacteria or photosynthetic eukaryotes, the genetic disruption of one FLV isozyme results in the loss of accumulation of the other at the protein level (Gerotto et al. 2016; Mustila et al. 2016), with the notable exception of the cyanobacterial FLV3 that is still present in the *flv1* deficient mutant (Mustila et al. 2016). In addition, the disruption of any *FLV* gene has a strong effect on O_2_ photoreduction in many environmental conditions (Chaux et al. 2017; Allahverdiyeva et al. 2011; Santana-Sanchez et al. 2019). However, the functioning of FLV as a hetero or homodimer remains blurry in most photosynthetic organisms, and the dissection of each hypothesis remains difficult. Here we have shown an atypical gene fusion of *FLVA* and *FLVB* in *Cladocopium* sp. that has an O_2_ photoreduction potential (Fig. 2 and 3). Gene fusion events are the result of strong selective pressure (Enright et al. 1999), thus making it clear that in the coral symbiont, both FLVA and FLVB are required for O_2_ photoreduction, at least, at the transcript level. Immunoblotting was likely to detect the fusion protein but also the individual proteins of FLVA and FLVB in *Cladocopium* sp. (Fig. 6), which suggests that the hybrid protein is cleaved by a post-translational mechanism. In the cyanobacterium S6803, the FLV3 homodimer does not function in O_2_ photo-reduction but in stress acclimation (Mustila et al. 2016). The fusion protein may be cleaved to form the homodimer in the Symbiodiniaceae *Cladocopium* sp. In photosynthetic eukaryotes like green microalgae, when transcriptomic data are available, the *FLVB* gene is much more expressed than the *FLVA* gene (Kleessen et al. 2015; Fang et al. 2012), which has long suggested a different role for each proteins formed (Mustila et al. 2016). In addition, transcriptomic data have found different expression patterns for both FLVA and FLVB genes (Fang et al. 2012; Tulin and Cross 2015), but the role of such differential regulation remains to be explored. Our results showing an atypical *FLVA*/*FLVB* polycistronic gene indicate that both FLV are probably required at the same quantity to be functional in coral symbionts.

Since the discovery, in the early 2000’s, a few groups have tried to measure the activities for recombinant photosynthetic FLV proteins, but almost negligible O_2_-reducing activities have been detected (Vicente et al. 2002; Shimakawa et al. 2015). Recently, some activities have been found in purified FLVA and FLVB homodimers from the cyanobacterium S6803 (Brown et al. 2019), but the catalytic turnover of 30 s^−1^ using NAD(P)H as the reductive agent seems to be still slow. However, our data clearly suggests that the presence of both proteins is crucial for FLV activity, we therefore propose that such *in vitro* activity of a homodimer is probably much lower than the one mediated by a heterodimer. The recently discovered NO-reduction mechanism of FLV (Burlacot et al. 2020) may be a new way to assess recombinant FLV activity with a more specific reaction. Recently, it has been implied that FLVA can be targeted by thioredoxin in *Chlamydomonas* (Pérez-Pérez et al. 2017), and actually several cysteine residues are conserved in both FLVA and FLVB sequences (Fig. 4). Redox regulation of FLV may also affect in vitro measurement of recombinant FLV and should be also studied further in future works to clearly define the O_2_ reducing activity.

Whereas genes encoding FLV are broadly conserved in algae and basal land plants in the photosynthetic green plastid lineage, Symbiodiniaceae is the only specie where a FLV can be found in the photosynthetic red plastid lineage until now. Interestingly, one big insertional *FLV* modification is conserved in *FLVA* only of *Chlamydomonas*, *Coccomyxa*, and basal land plants (Supplemental Fig. S1), which suggests that this modification occurred during the evolutionary history of photosynthetic green plastid lineage. That is, the Symbiodiniaceae FLV have evolved through the different evolutionary history from the green plastid lineage. The phylogenetic tree suggested that FLV in Symbiodiniaceae is highly likely to have been inherited from basal green algae (Fig. 5). One possibility is that the dinoflagellates obtained the genes for FLVA and FLVB in the polycistronic form in a lateral gene transfer event (Fig. 7). As the other possibility, the FLV genes might be inherited from basal green algae by the transient endosymbiosis (Keeling 2013). We cannot exclude the possibility that Symbiodiniaceae has obtained FLV at the secondary endosymbiosis with basal green algae (Fig. 7), although dinoflagellates are often recognized as secondary algae derived from red algae (Falkowski et al. 2004). Ultimately, it is possible that *FLV* genes had been obtained from basal red algae at the secondary endosymbiosis before they have been lost in photosynthetic red plastid lineage, although there is no other report showing red algae harboring FLV. Recently, we found that P700 oxidation depends on the presence of O_2_ in several secondary algae in the photosynthetic red plastid lineage *in vivo*, like in cyanobacteria and green algae (Shimakawa et al. 2019). Whether these O_2_-dependent P700 oxidation could be linked to FLV inherited from cyanobacteria or basal green algae remains to be explored in the future.

**Fig. 7.**
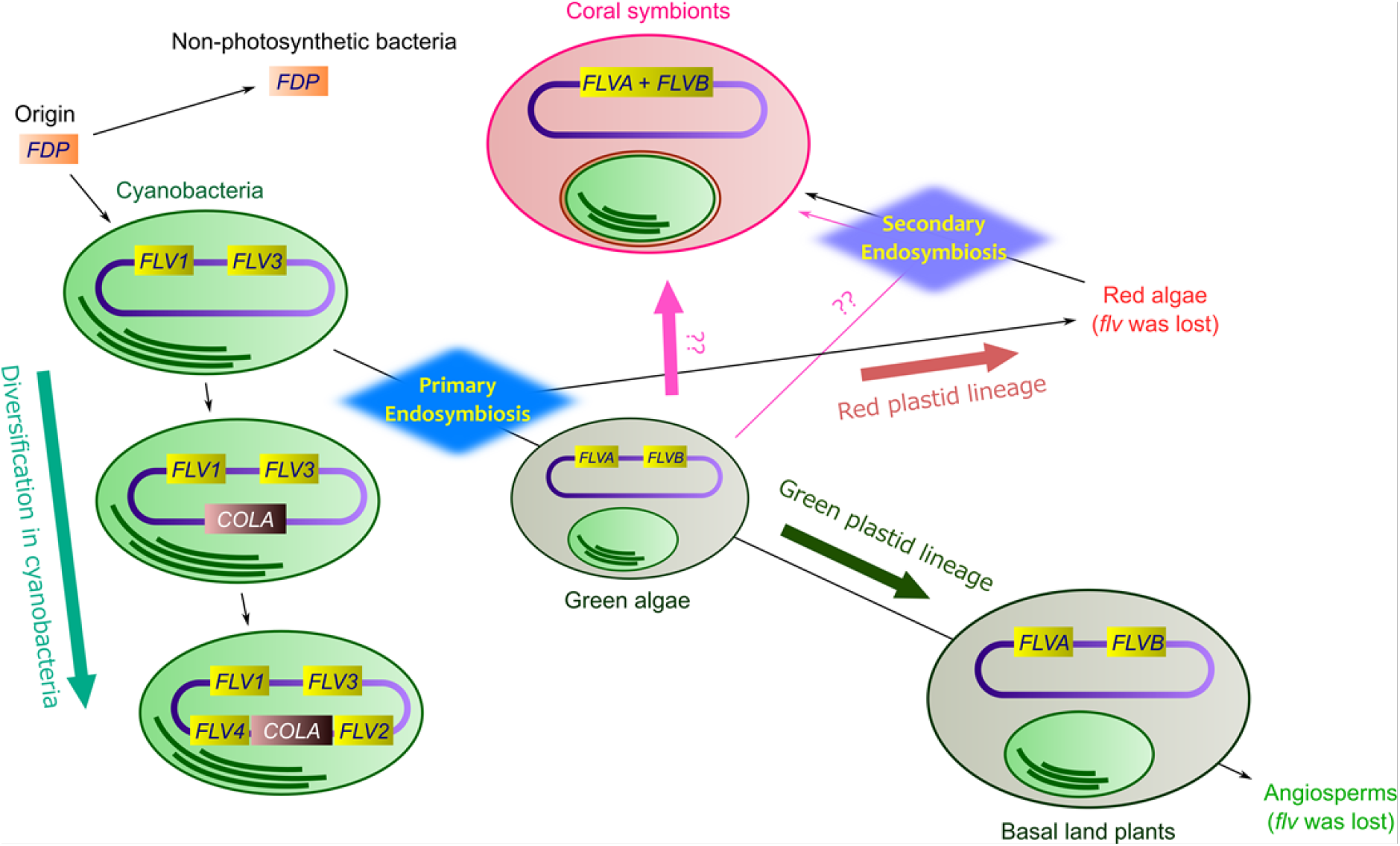
A hypothetical model for the epidemic events of genes encoding flavodiiron proteins (FLV) in photosynthetic organisms. Originally, FLV is categorized into one group of the protein family (more generally termed as FDP). Gene duplication have generated two types of *FLV* genes in cyanobacteria, which have been inherited to green algae but lost in photosynthetic red plastid lineage. Some cyanobacteria species had obtained a CO_2_-limitation associated protein (*COLA*) and sequentially constructed *flv4-2* operon. In the coral symbiont Symbiodiniaceae, *FLV* genes are assumed to have been inherited from basal green algae.

There are some cases of gene fusion events reported in dinoflagellates. For example, a paralogue of 3-dehydroquinate synthase (a shikimate biosynthetic enzyme) and a caffeoyl CoA 3-*O*-methyltransferase have been inherited from cyanobacteria to the dinoflagellate *Oxyrrhis marina* by lateral gene transfer, resulting in the fusion protein (Waller et al. 2006). Additionally, a glyceraldehyde 3-phosphate dehydrogenase and an enolase are also encoded in the form of gene fusion in the dinoflagellates *Karenia* and *Heterocapsa* (Takishita et al. 2005). Further, up to 10 small subunits of the light-harvesting complex are expressed in the form of one fusion protein, and then the polypeptide is cleaved presumably at each arginine position after translation in the dinoflagellate *Amphidinium carterae* (Hiller et al. 1995). Whereas it has been accepted that dinoflagellates have a different approach of transcription and gene expression compared with other photosynthetic organisms (Bachvaroff and Place 2008), the molecular mechanism of these atypical post-translational cleavage of fusion protein is still poorly understood. In the case of Symbiodiniaceae FLV, the specific insertional modification at FlR-like domain in the upstream region (Supplemental Fig. S1) may function in being recognized by a post-translational cleavage system. Based on the amino acid sequences of each upstream and downstream part, individual FLVA and FLVB in Symbiodiniaceae were both predicted to be localized in chloroplasts except for the upstream region of FLV in *Symbiodinium tridacnidorum* (Supplemental Table S1). Whether the fusion protein may be cleaved before the translocation from nucleus to chloroplasts is still to be clarified.

It is astonishing that coral symbionts are the only algae found to harbor FLV in the photosynthetic red plastid lineage. The possible explanations could be linked to the symbiotic way of living of the coral symbionts. Since symbionts are fixed inside of the coral reefs, they cannot rely of distribution across the water depth in the shallow areas in the sea to protect from excess light energy and require strong and resilient photoprotection mechanisms like P700 oxidation mechanism for protecting PSI against photodamage. Another possibility is that since the FLV-mediated O_2_ photoreduction creates lumen acidification by driving photosynthetic linear electron flow, it contributes to the additional ATP production and the increase in stromal pH that could favor a locally outside pH decrease, promoting calcification. The physiological role of such exceptional FLV that catalyzes O_2_ photoreduction in the coral symbiont remains to be investigated in future works and may give us new insights on how corals will respond to the forthcoming environmental changes.

## Conflict of interest

The authors have no conflict of interest to declare.

## Accession numbers

Nucleotide and amino acid sequences data studied in this study can be found on the database of National Center for Biotechnology Information (NCBI, https://www.ncbi.nlm.nih.gov), JGI Genome Portal (https://genome.jgi.doe.gov), OIST Marine Genomics Unit (https://marinegenomics.oist.jp), CyanoBase (http://genome.microbedb.jp/cyanobase), and MarpolBase (https://marchantia.info) following the accession numbers: *Escherichia* FDP (WP_000029589), S6803 FLV1 (Sll1521), FLV2 (Sll0219), FLV3 (Sll0550), and FLV4 (Sll0217), S7942 FLV1 (1810) and FLV3 (1809), S7002 FLV1 (A1743) and FLV3 (A1321), *Chlamydomonas* FLVA (Cre12.g531900) and FLVB (Cre16.g691800), *Coccomyxa* FLVA (54807) and FLVB (45048), *Micromonas* FLVA (90715) and FLVB (58403), *Ostreococcus* FLVA (e_gw1.02.00.129) and FLVB (fgenesh1_pm.C_Chr_02.0001000066), *Marchantia* FLVA (Mapoly0005s0210) and FLVB (Mapoly0103s0039), *Physcomitrella* FLVA (Pp3c14_14450V3) and FLVB (Pp3c1_26720V3), *Selaginella* FLVA (143660) and FLVB (448328), *Symbiodinium* FLV (s632_g30), *Cladocopium* FLV (comp44467), and *Durusdinium* FLV (TRINITY_DN39309).

## Acknowledgements

The authors gratefully thank Dr. Anja Krieger-Liszkay (CNRS Paris-Saclay, France) for the kind advices in writing the manuscript. The authors thank Dr. Gilles Peltier (CEA Cadarache, France) for kindly providing FLVA antibody.

## Notes

**Funding information:** This work was supported by the Japan Society for the Promotion of Science (JSPS; grant no. 20J00105 to G.S.).

### Competing Interest Statement

The authors have declared no competing interest.

