## Supplementary material for "Coral symbionts exhibit a polycistronic flavodiiron gene leading to functional proteins in photosynthesis": SupplementalFigS1_CoralFlv_20210403.pdf

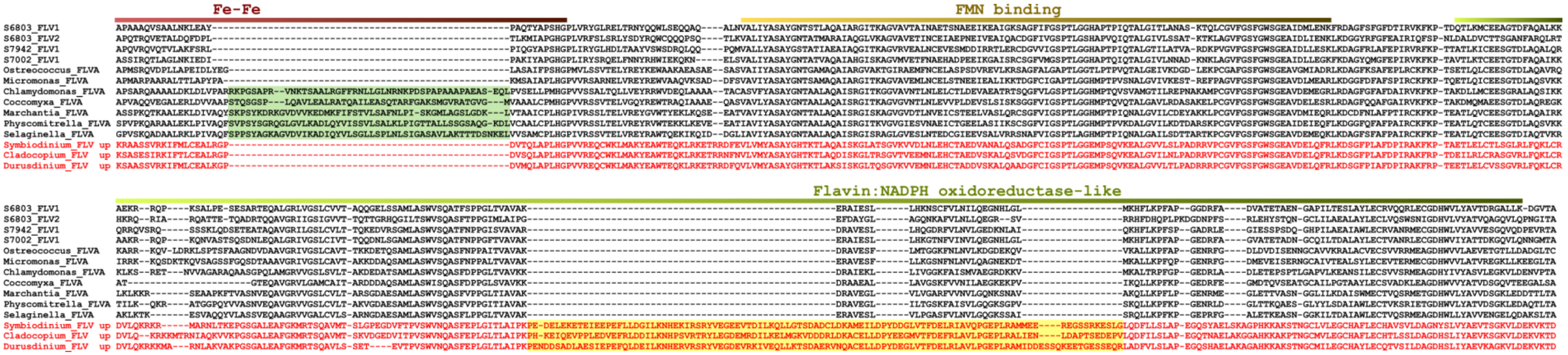


**Supplemental Fig. S1.** Amino acid sequence alignment of flavodiiron proteins A (FLVA) in cyanobacteria *Synechocystis* sp. PCC 6803 (S6803), *Synechococcus elongatus* PCC 7942 (S7942), and *Synechococcus* sp. PCC 7002 (S7002), green algae *Ostreococcus tauri*, *Micromonas* sp. RCC299, *Chlamydomonas reinhardtii*, and *Coccomyxa subellipsoidea*, the liverwort *Marchantia polymorpha*, the moss *Physcomitrella patens*, the fern *Selaginella moellendorffii*, and dinoflagellates Symbiodiniaceae (coral symbionts) clade A3 (*Symbiodinium* sp.), C (*Cladocopium* sp.) and D (*Durusdinium* sp.). The sequences in Symbiodiniaceae were analysed in the separated form to upstream (up) region. Brown, yellow, and green bars indicate diiron centre (Fe-Fe), flavin mononucleotide (FMN) binding domain, and flavin:NAD(P)H oxidoreductase-like motif, respectively. Green and yellow shadings present the insertional sequences specific to the photosynthetic green plastid lineage and to Symbiodiniaceae.
