## Supplementary material for "Coral symbionts exhibit a polycistronic flavodiiron gene leading to functional proteins in photosynthesis": SupplementalTableS1_FusionFLV_20210330.pdf

**Supplemental Table S1.** Predicted subcellular localization (%)

| Species | Chloroplast | Mitochondrion | Cytoplasm | Others |
|---|---|---|---|---|
| *Symbiodinium tridacnidorum* | | | | |
| Upstream | 15.0 | 82.5 | 1.2 | 1.3 |
| Downstream | 87.7 | 11.6 | 0.3 | 0.4 |
| *Cladocopium* sp. | | | | |
| Upstream | 97.8 | 0.7 | 0.3 | 1.2 |
| Downstream | 72.8 | 13.8 | 8.6 | 4.8 |
| *Durusdinium trenchii* | | | | |
| Upstream | 64.8 | 12.6 | 13.8 | 8.8 |
| Downstream | 53.3 | 46.4 | 0.1 | 0.2 |
